# Label-Free Longitudinal Imaging of Single Cell Drug Response with a 3D-Printed Cell Culture Platform

**DOI:** 10.1101/2025.08.02.668298

**Authors:** Erin L. Dunnington, King Wai Chiu, Brian S. Wong, Anthony Chales-Antonio, Victoria Pang, Yiyi Wu, Dan Fu

## Abstract

Image-based phenotypic screening has emerged as a powerful tool for revealing single-cell heterogeneity and dynamic phenotypic responses in preclinical drug discovery. Compared to traditional static end-point assays, live-cell longitudinal imaging captures the temporal trajectories of individual cells, including transient morphological adaptations, motility shifts, and divergent subpopulation behaviors, enabling high content features and more robust early prediction of treatment outcomes. Fluorescence-based screening, while highly specific, is constrained in live-cell contexts by broad spectral overlaps (limiting multiplexing to fewer than six channels), bulky fluorophores that may perturb small-molecule interactions, and photobleaching or phototoxicity under repeated excitation. Stimulated Raman scattering (SRS) microscopy overcomes these barriers by delivering label-free, quantitative chemical contrasts alongside morphological information. Here, we present a low-cost, 3D printed cell culture platform compatible with the stringent optical requirements of SRS microscopy. This set up enables real-time drug delivery and continuous monitoring of biochemical and morphological changes in living cells during 24-hour time-lapse imaging with minimal photodamage. We outline a processing pipeline for longitudinal SRS images to extract chemical and morphological features of single live cells. Using this system, we showcase time-lapse SRS microscopy as a tool to map heterogenous drug-induced single-cell response over time, enabling the identification of varying trajectories within complex cell populations. By parallelizing multi-well perfusion with label-free chemical imaging, our approach offers a pathway toward high-throughput pharmacodynamic assays for the acceleration of phenotypic screening and personalized medicine.

## INTRODUCTION

The progression of microscopy as an effective cell response screening tool has been critical in advancing the preclinical drug discovery and development process. While traditional cell-based assays (e.g. MTT, ATP luciferase bioluminescence) provide rapid readouts of cell health and function, image-based cytometry is gaining prominence as a means to quantitatively uncover drug- or genetic modification-induced phenotype changes in single cells.^1–5^ These imaging methods offer a detailed view of cellular heterogeneity, including identifying response subpopulations and drug mechanism of action. Phenotype response screening generally utilizes optical images to generate rich datasets that necessitate the use of specialized analysis software or machine learning algorithms to link observed features with disease-relevant phenotypes.^6^ The addition of live cell tracking allows for measurements of cell motility and changes in morphology and signal intensity that can further reveal heterogeneous cellular behavior beyond that in a single snapshot.^7–9^

Traditionally, fluorescence-based screening has been the workhorse for live-cell assays. It combines automated multiwell plate imaging with the use of light-emitting chemical probes to visualize different cell types, subcellular structures, and biomolecules. Despite its specificity, long-term fluorescence imaging is often constrained by the overlap of broad spectra, which typically limits the number of imaging channels to no more than six colors.^10, 11^ Furthermore, most drug molecules cannot be directly visualized with fluorescence. Fluorescent labels are often larger than small-molecule drugs, which can alter their interactions with cellular targets.^12, 13^ Fluorophores also suffer from photobleaching or phototoxicity issues after periodic excitations.^13^ Label-free optical imaging can partly address the limitations of fluorescence imaging of live cells. For example, brightfield microscopy can been paired with machine learning to distinguish drug-induced cell morphological variations.^14^ Differential interference contrast (DIC)^15^ and quantitative phase imaging (QPI)^8, 16^ can reveal longitudinal cell morphology and mass dynamics by mapping refractive index difference or change in optical path length.^8, 17^ However, these label-free optical techniques lack the chemical specificity and organelle specific contrast seen in fluorescence microscopy, which are often necessary to monitor molecular events such as metabolic shifts or drug-cell interactions. Spontaneous Raman microscopy offers molecular-specific contrast but requires integration times on the order of seconds per spectrum, which makes live cell imaging impractical.^18^

Stimulated Raman scattering (SRS) microscopy is a high-resolution, label-free chemical imaging technique that generates images with molecular vibrational contrast at high speed up to video-rate.^19, 20^ Importantly, multiple narrow spectral SRS bands can be quickly captured to resolve different molecular species. SRS signal intensity is linearly dependent on the concentration of bonds, allowing for quantification of chemical content.^21^ Despite these advantages for phenotypic screening, long-term SRS imaging of live cells has been scarce. This is due to the incompatibility of most bulky commercial stage-top incubators with the short working distances of high-numerical aperture (NA) objective and condenser lenses used for SRS imaging. Kim et al. addressed this by presenting a thin microfluidic chip with integrated valves fabricated with PDMS-based soft lithography^22^. Ota et al. combined Raman and SRS with a PDMS microfluidic chip to observe dynamic paramylon production in *E. gracilis* under controlled flow at two timepoints.^23^ Others have integrated similar devices with environmental control for longitudinal imaging. For example, Yuan et al. fabricated a thin, flexible silicon rubber chamber for upright SRS setups, enabling 24-hour time-lapse cell imaging with minimal phototoxicity.^24^ Relatedly, Tipping et al. utilized a commercial perfusion chamber with pipetting to wash and label cells for long-term monitoring of the uptake and distribution of the DDR1 inhibitor 7RH.^25^ Tague et al. employed longitudinal SRS imaging to monitor fatty acid accumulation across the same population of single *E. coli* on agarose pads kept at 31°C.^26^ Stiebing et al. traced deuterated palmitate uptake in live human macrophages long-term.^27^ However, to our knowledge, no existing platform integrates environmental control with multichannel perfusion chambers optimized for SRS’s stringent optical and detection requirements, leaving a critical need for systems that can parallelize drug dosing and chemical imaging under physiologically relevant conditions. Likewise, we are unaware of any published study that utilizes segmentation and tracking to extract single cell feature trajectories from longitudinal SRS images.

Here, we present a 3D-printed stage-top cell culture platform built around an inverted SRS microscope that enables multi-channel on-demand drug delivery and continuous monitoring of biochemical and morphological changes of single cells. The device is readily produced with a consumer 3D printer at low cost. It contains multiple independent flow channels with precise temperature control that preserve cell viability and mimic physiological conditions. During 24-hour SRS imaging, we observe little photodamage-related effects exhibited in live cells. We outline a processing pipeline for longitudinal SRS images to extract and quantify key morphological and chemical features of single-cell responses to diverse anticancer agents. Using this system, we showcase longitudinal SRS microscopy as a tool to map dynamic single-cell heterogeneity over time, enabling the identification of varying trajectories within complex cell populations and classification of different drug mechanisms. By integrating parallel perfusion control with chemical imaging, our device enables high-content phenotypic evaluation of multiple treatments simultaneously in a single experiment. This work represents a major step towards applying label-free chemical imaging towards phenotypical drug screening. The unique ability of SRS microscopy to quantify both metabolic shifts and intracellular drug uptake in 2D and 3D cultures^28–30^ suggests that our cell culture and imaging platform could be ideally suited for high-content patient-derived tissue or organoid assays with time-resolved pharmacodynamic information, promising deeper mechanistic insights and a path toward personalized medicine.

## MATERIALS AND METHODS

### Device fabrication

3D models of the device were created using Autodesk Fusion 360. The perfusion chambers were fabricated out of polylactic acid (PLA) using a P1P 3D printer (Bambu Lab) equipped with a 0.2 mm hot end. After printing, the device was sterilized with 75% ethanol and UV light for 15 minutes. 25×60 mm coverslips were affixed to the top and bottom of each chamber autoclaved 19-gauge stainless steel tubing with 90° bend (McMaster-Carr) were secured with a thin layer of polydimethylsiloxane (PDMS, Dow Chemical Company) as the inlets and outlets of the chamber. PDMS was prepared according to the manufacturer’s instructions and cured at room temperature for 72 hours. The device was thoroughly rinsed with sterile PBS prior to cell seeding.

### Cell culture

The human lung adenocarcinoma A549 cell line (ATCC) was maintained at 37°C in a 5% CO_2_ (*v/v*) humidified incubator. A549 cells were cultured in Dulbecco’s modified eagle medium (DMEM, Gibco) supplemented with 10% fetal bovine serum (Hyclone) and 1% penicillin-streptomycin (Gibco). A549 cells were seeded into the device at a density of 2 × 10^4^ cells/well by syringe and incubated for 24 hours prior to imaging. An additional 15 mM HEPES buffer was added to the culture medium to increase the medium buffering capacity.^31^

### Drug treatment

Osimertinib (SelleckChem), lapatinib (SelleckChem), paclitaxel (Sigma-Aldrich) and doxorubicin (Sigma-Aldrich) were dissolved in dimethyl sulfoxide (DMSO) to a 10 mg/mL stock solution. DMSO-treated cells were used as vehicle control. Drugs were diluted to the desired concentration in DMEM cell culture medium.

### CellTiter 96® AQueous Cell Proliferation Assay

A549 cells were seeded at a density of 4 × 10^4^ cells/well in 96-well plates for 24 hours prior to 24-hour incubation with drugs. Following treatment, cells were incubated with serum-free media and combined MTS/phenazine methosulfate solution (Promega) for 1 hour. Absorbance measurements were collected at 490 nm using a UV/visible microplate reader (Thermo Labsystems Multiskan Spectrum).

### Device heating and temperature regulation

A 1.1 mm thick optically transparent indium titanium oxide (ITO) glass slide (30-60Ω) (Delta Technologies) was placed on top of the device for heating. Copper wires were soldered onto copper tape on the conductive side of the ITO slide to connect to an autotunable Proportional-Integral-Derivative (PID) temperature controller (CN38S, Omega Engineering). A 10kΩ negative temperature coefficient (NTC) thermistor (Newark Electronics) was inserted into a chamber to provide readouts back to the PID loop. The ITO slide was pre-heated prior to placement on the 3D print to avoid exposing the sample to harsh temperature changes. For tracking temperature stability in Figure 1, an additional sensor (Okolab) was inserted into a separate well in the chamber for temperature recording.

**Figure 1.**
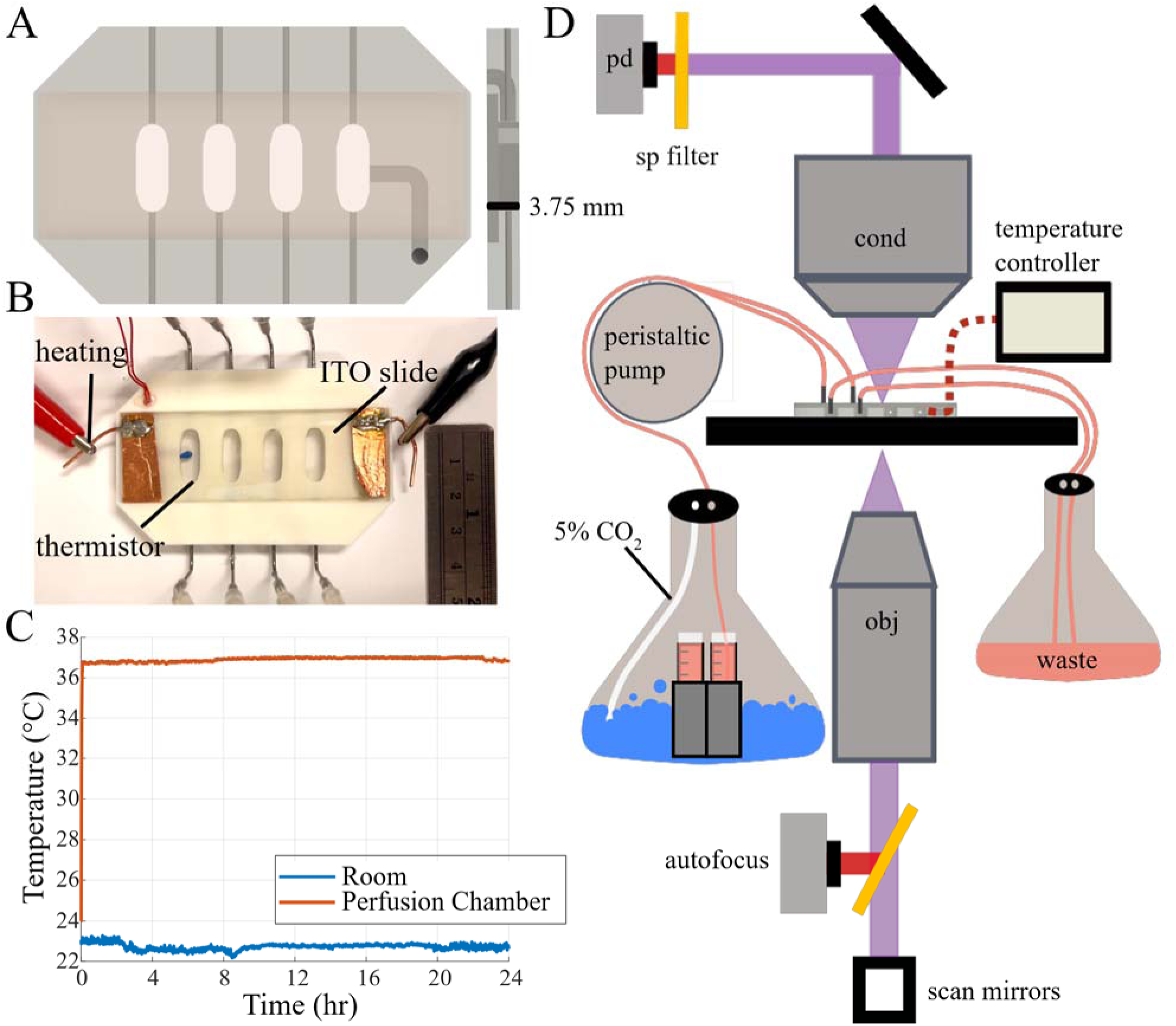
SRS microscope with live cell imaging compatibility using a 3D-printed perfusion chamber. A. 3D-rendered representation of the top-view and side-view of device with labelled device height. B. Image of perfusion chamber with addition of tubing and heating components. A ruler is provided for size reference. C. Trace of internal chamber temperature with perfusion against room temperature over 24 hours. A second temperature sensor was placed alongside the thermistor in the same experimental conditions as on-device culture for temperature recording. Measurements were recorded every 10 s immediately after the onset of perfusion. D. Schematic diagram of SRS microscope setup highlighting the imaging and cell culture components. PD: photodiode; BPF: bandpass filter; Cond: microscope condenser; Obj: microscope objective; DCM: dichroic mirror.

### Perfusion control

A peristaltic pump (L100-1S-1, Longer Precision Pump Company) was used to control the perfusion of cell culture medium through the chambers using soft clear Versilon PVC tubing (1/16” ID, 1/8” OD) with a FEP (fluorinated ethylene propylene) lining (6519T16, McMaster-Carr). FEP is a fluoropolymer with high chemical resistance, low adsorption, and low gas permeability.^32, 33^ The outer layer of flexible polymer provides the mechanical strength and flexibility necessary for peristaltic pumping. The outer layer of flexible polymer provides the mechanical strength and flexibility necessary for peristaltic pumping. Cell culture medium was pre-loaded into the tubing and continually drawn from the medium reservoir through the device at a flow rate of 2.0 µL/min. As illustrated in Figure 1D, 5% CO_2_ dissolved in air is bubbled into sterile water at the bottom of a flask and equilibrates into drug-dosed cell culture medium.

### Cell counting

Images were collected with a brightfield microscope (CK2, Olympus) equipped with a 20x, 0.4 NA air objective (LWD C A 160/1.2 A20, Olympus). To capture the same field of view, the bottoms of each well were marked with a glass etcher at five different locations. The seeding density in the six-well plate was equivalent to that in the perfusion chamber by accounting for the surface area, with an equivalent scale up in medium volume.

### Longitudinal SRS imaging

SRS images were acquired using a home-built microscope as described in our previous publications.^34^ A femtosecond dual-beam Insight DS+ (Spectra-Physics) laser outputs a fixed beam at 1040 nm (Stokes) and a tunable beam (pump) set at 798 nm with an 80 MHz repetition rate. The 1040 nm Stokes beam is then modulated at 20 MHz by an electro-optical modulator (EOM, Thorlabs) and coupled into a 4-m polarization-maintaining Yb-doped fiber (YB1200-10/125 DC-PM, Thorlabs) for parabolic amplification.^35^ Pulse chirp is controlled using 64 cm of high-dispersion dense flint glass rods (Newlight Photonics) and a grating stretcher (LightSmyth) for the pump and Stokes beams, respectively. The two beams are combined with a dichroic mirror and temporally overlapped using a motorized Zaber X-DMQ-AE delay stage. They are then directed into an inverted laser scanning microscope (IX73, Olympus) equipped with a 20x, 0.75 NA air objective (UPLSAPOP20X, Olympus) and a 0.8 NA air condenser (FN-C LWD, Nikon). The pump and Stokes beams were both set to 40 and 50 mW at focus, respectively. After the condenser, the pump beam is isolated with a 1000 nm shortpass filter (SP, ThorLabs) and detected by an amplified Si photodiode (PD, Hamamatsu). The signal is then demodulated with a lock-in amplifier (H2FLI, Zurich Instruments) with a time constant of 4 µs. A Continuous Reflection Interface Sampling and Positioning (CRISP) autofocus system (Applied Scientific Instrumentation, ASI) is mounted at the camera port of the microscope and connected to an automated MS-2000 *XYZ* stage controller for focus locking. A 700 nm short pass dichroic mirror is used to reflect the pump and Stokes beams while transmitting the 650 nm LED required for focus locking. All images collected were 1024 × 1024 pixels with a pixel size of 0.33 μm. Three-band images at 2850 cm^-1^ (–CH_2_ stretching, lipid), 2930 cm^-1^ (–CH_3_ stretching, protein) and 3170 cm^-1^ (O–H shoulder, water) were collected for each field of view every 20 minutes for all live-cell experiments. Image capture through ScanImage Basic software (MBF Science) and sample focus across the multi-well device were controlled *via* a custom Matlab script.

### Cell and vacuole segmentation

After time-lapse image collection, each frame was field normalized. Single cells from the normalized SRS image at each time point were segmented with the deep learning-based Cellpose-SAM model that was retrained with manual segmentation.^36^ Persistent errors in the segmentation mask were manually corrected in ImageJ. A separate Cellpose-SAM model was individually retrained for vacuole segmentation across all drug treatments, primarily Lap and Osi.

### Cell tracking

Cell segmentation masks were used for single-cell tracking in TrackMate^37^, an ImageJ/Fiji plugin. The label image detector and Advanced Kalman Tracker were utilized. Tracks totaling ≤ 20 frames and tracks of cells on the edge of the field of view were removed from further analysis. A custom Python script was utilized to rebuild the lineage tree from TrackMate for correlation analysis with extracted features.

### Feature extraction

With a binary cell segmentation mask, cell intensity statistics were extracted using SciPy.^38^ Scikit-image provides classical shape descriptors, moments, and inertia tensor measurements based on a binary mask alone.^39^ Haralick textures were computed by mahotas.^40^ Centroid coordinates of cells were extracted for cell motility analysis. Quantification of vacuole statistics were done by using binary vacuole mask and cell mask. All 167 features were collected in Python, and corresponding details are listed in Table S1.

### Statistical analysis

For each endpoint measurement, including foldchange in cell count, time-resolved feature ratios, and vacuolization metrics, differences among treatment groups were evaluated by one way analysis of variance (ANOVA). Where the ANOVA indicated a significant overall effect (α = 0.05), pairwise comparisons were made using Tukey’s post-hoc test. Statistical significance is denoted as *p < 0.05, **p < 0.01, and ***p < 0.001. All data were analyzed in MATLAB R2024b (MathWorks, Natick, MA). For all plots single-cell measurement plots containing an error bar, this indicates interquartile range (IQR), unless started otherwise.

## RESULTS AND DISCUSSION

### Live-cell perfusion chamber design and characterization

A low-cost cell culture perfusion chamber was integrated into our homebuilt inverted SRS laser scanning microscope system for optimal on-stage time-lapse imaging of live adherent cells. The device (Figure 1A) was fabricated out of PLA, a low-cost 3D printing material often used to mold scaffolding and channels in microfluidic and organ-on-a-chip *in vitro* devices.^41, 42^ PLA was chosen due to its biocompatibility and inability to absorb small molecules such as chemotherapeutics.^43^ Briefly, the device is composed of four wells, one for temperature control while under perfusion and three for cell culture and imaging. The chamber is created by sandwiching the PLA print with two standard #1.5 glass coverslips. L-shaped metal tubes are used as inlets and outlets for perfusion. Each well has a surface area of ∼87 mm^2^, for a total culture well volume of 326 µL. As indicated in the side view, the whole chamber is constrained to 3.75 mm tall to fit within the working distance of the objective and condenser used, approximately 7 mm.

Live cell culture requires tight regulation of several parameters, including temperature, osmolarity, humidity, pH/CO_2_ concentration, and nutrients supply. It is important to ensure that the cells are responding to the chemicals introduced and not to uncontrolled environmental conditions in drug screening assay.^44^ The temperature should be maintained near 37°C for optimal growth and physiology of mammalian cells.^45^ For our perfusion chamber, optically transparent ITO-coated glass slides were used to heat the sample (Figure 1B). Due to its low electrical resistance, ITO-glass substrate generates heat when an electric current passes through it. ITO glass-based heaters have previously been used for microfluidic cell culture chips.^46^ A tuned PID loop is used to maintain a consistent temperature at the setpoint. After 5 minutes, temperature stabilization near 37.0°C was maintained over the course of 24 hours, with a total fluctuation between 36.69°C and 37.06° (Figure 1C). Comparatively, the room temperature fluctuated between 22.17°C and 23.28°C at the same time. At physiological temperatures, cell culture medium can evaporate and disrupt water balance that can lead to osmotic stress responses and increased drug concentrations.^45, 47^ With the use of ITO heating in close proximity to the sample, we found it difficult to minimize evaporation in an open design system. When mineral oil is used to reduce evaporation, it gives rise to a shifting SRS background due to a lens effect from the uneven mineral oil layer (Figure S1). Medium evaporation is eliminated in the closed system, creating a stable microenvironment that allows for perfusion of fresh media, ensuring consistent osmolarity.

A slow, continuous flow of CO_2_ equilibrated medium through the device assists in supplying fresh media to the cells to maintain nutrient supply and pH and remove waste. The regulation of these components is essential for healthy cell culture: 1. pH influences enzyme activity, membrane stability, and metabolic processes and is regulated by equilibration of CO_2_ with bicarbonate-buffered media; 2. Continuous supply of nutrients support energy production and biosynthesis; 3. Efficient waste removal prevents the buildup of toxic byproducts of cellular respiration and metabolism that can compromise cell viability.^48, 49^ The flow through each channel in the chamber is visualized in Movie S1. A full list of device supplies and their costs is provided in Table S2.

Once assembled, the perfusion chamber is secured on an inverted laser-scanning SRS microscope. As the device is heated, focus drift due to temperature fluctuations must be addressed to maintain high resolution imaging. A hardware autofocus system was utilized to actively detect and correct changes in the distance between the objective lens and the sample in real time. We used a moderate numerical aperture air objective and condenser to eliminate the need for immersion medium such as oil or water. This reduces the risk of evaporation or additional sample disturbance and focus drifting over extended time-lapse experiments. It also increases the field of view for observation due to lower magnification without significantly degrading spatial resolution.

### Evaluation of on-device live cell viability with continuous SRS imaging

After device optimization, live A549 cell viability within the perfusion chamber was evaluated by measuring the fold change in cell count over 24 hr. First, on-device growth was compared to a traditional CO₂ incubator. Under our culture conditions, the vendor reports a characteristic doubling time of ∼22 h, equivalent to an expected ∼2-fold increase in cell number over 24 hr. In proliferating mammalian cells, biosynthetic flux, cell cycle duration, and growth rate are coregulated to preserve homeostasis; thus, maintenance of this ∼2-fold change serves as a proxy for healthy, noncytotoxic conditions.⁴⁶ Cell counts were obtained from brightfield images of fields of views starting ≥200 cells each.

In a standard, plasma-treated 6-well plate, A549 cells exhibited a 2.02--fold increase (±0.32) in cell number over 24 h (Figure 2A). In our 3D--printed chamber housed inside the CO₂ incubator, the fold change was 1.76 (±0.40). Following transfer to the onstage perfusion setup, where fresh medium was supplied at 2.0 µL/min (device refresh ≈3.75 hr) to minimize metabolite accumulation and shear stress (∼2.3 × 10⁻⁵ dyn- cm⁻²), the fold change measured was 1.83 (±0.30). Any differences in growth rate in the well-plate and 3D-print likely arise from increased cell attachment affinity for the plastic substrate of the well-plate over the glass coverslip. However, no statistically significant differences were found among these conditions, indicating that neither confinement in the device nor the addition of continuous perfusion impaired normal proliferative capacity.

**Figure 2.**
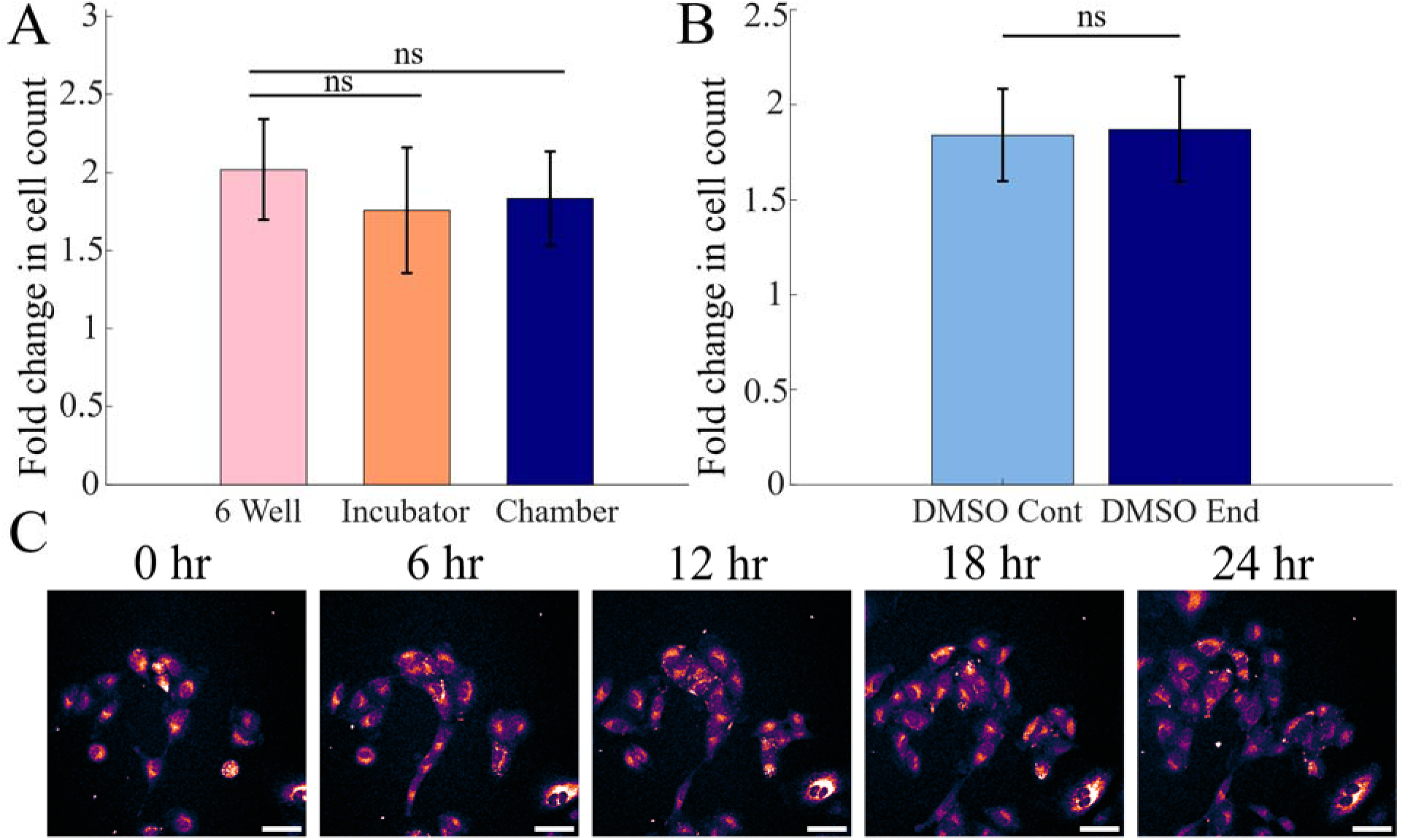
Evaluation of live cell viability in perfusion chamber. A. Fold-change in cell count over 24 h in a six-well plate (N = 10 FOV), 3D-printed chamber inside a CO₂ incubator (N = 10 FOV), and 3D-printed chamber with perfusion (N = 10 FOV). B. Fold-change in cell count when cells were imaged continuously (N = 8 FOV) versus only at the start and end of the session (N = 10 FOV). C. Representative SRS intensity projections of combined protein and lipid bands in DMSO-treated A549 cells imaged every 20 min. Scale bar: 50 µm. ns: not significant.

Previous studies have shown that time-lapse imaging is feasible with femtosecond pulsed laser-based imaging modalities as certain cell lines have strong tolerance to near-IR laser exposure over a wide range of imaging power and intensity.^26, 50, 51^ To assess phototoxicity of continuous SRS imaging, we compared fold change in cell count when imaging every 20 min (1.84 ± 0.24) to counts obtained only at the beginning and end of a 24 hr session (1.87 ±0.27) (Figure 2B). No significant difference was observed between the two regimens, demonstrating that our moderate NA air objective and optimized laser power minimize photodamage. Moreover, SRS images of protein and lipid bands acquired every six hours showed no increase in lipid droplet accumulation, an indicator of cellular stress, under continuous imaging (Figure 2C).

### Quantification of single-cell spectral and morphological features from longitudinal SRS images

We imaged A549 cells dosed with a range of anticancer treatments over time to evaluate the ability of longitudinal SRS imaging to examine drug-induced phenotypic changes in single cells. Per device, three wells were imaged: a vehicle control and two drug-treated conditions. For each condition, 73 images at 20 minutes intervals were captured for eight different fields of view. Each image consisted of three SRS spectral bands: lipid, protein, and water. For single-cell analysis, the lipid and protein frames of each image were intensity-averaged to increase contrast for segmentation using Cellpose-SAM (Figure 3A, B).

**Figure 3.**
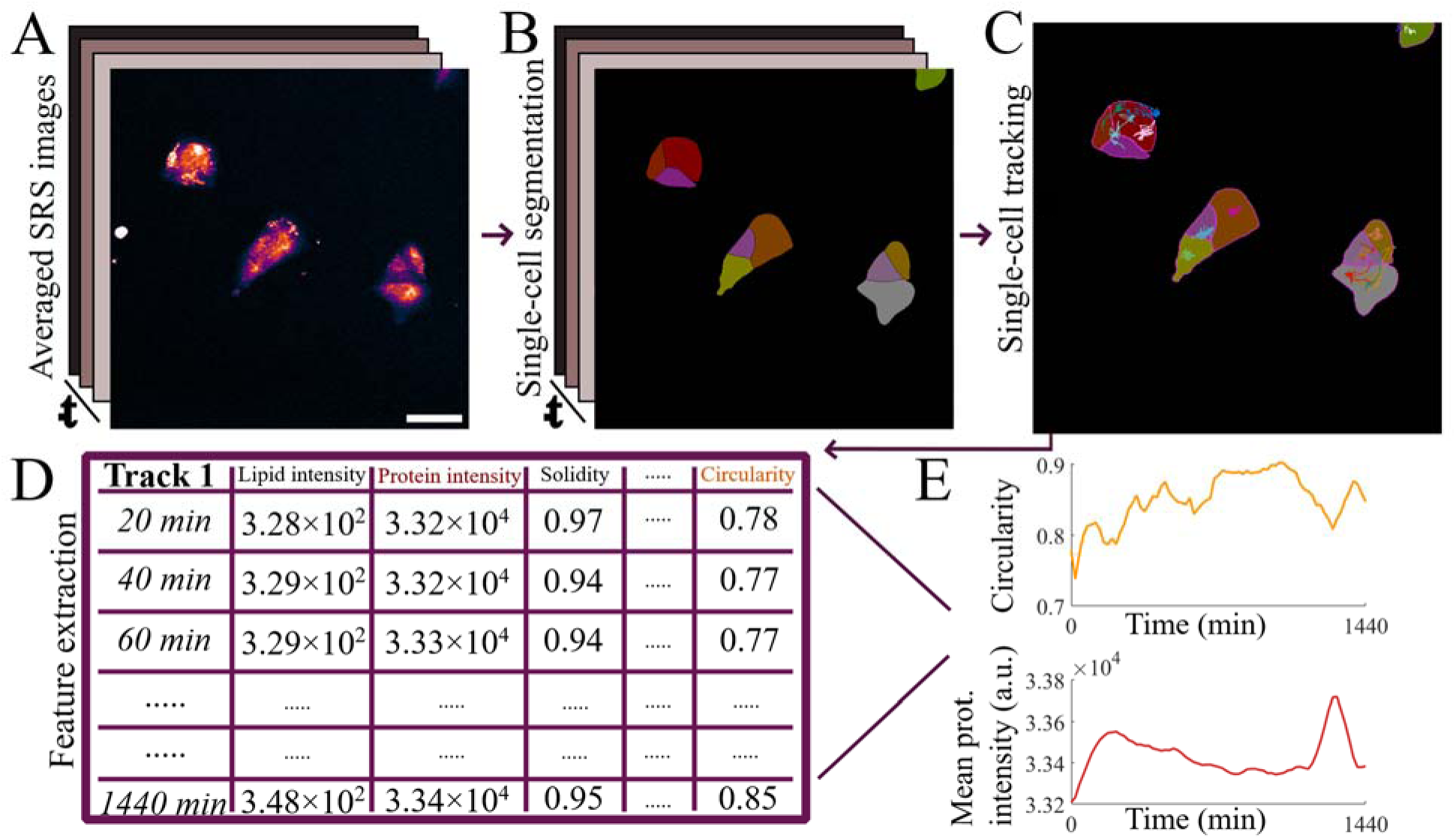
Single-cell feature extraction from 3-band longitudinal SRS images. A. Averaged intensity protein/lipid SRS images of Osi-treated cells over time. Scale bar: 50 µm. B. A mask is created from segmentation model, where each color is a different cell. C. Single-cell lineages are tracked from the segmentation mask. D. Cellular features are extracted from each frame and connected throughout temporal images using the tracked lineage tree. E. Features such as circularity and mean protein signal intensity of one cell can be observed over time.

A segmentation mask was generated for each of the 73 images in a stack, allowing for subsequent manual corrections to be performed in ImageJ. Following segmentation, each whole individual cell was detected throughout the stacked segmentation mask using TrackMate (Figure 3C). The Advanced Kalman tracker was found to be most suitable for our images due to the ability to detect track fusion and track division. The output of TrackMate allowed for the reconstruction of the lineage tree of each track. With this, every individual cell and division/fusion event could be identified (Figure S2).

Following tracking, morphology- and intensity-based (mean, standard deviation, min, max, skewness, and kurtosis) features of each single cell were extracted using an in-house script (Figure 3D, 3E). The regions of interest (ROIs) from the tracked segmentation masks were applied to the individual field-normalized SRS images of lipid, protein, and water for intensity-based feature extraction, while an intensity average of the lipid and protein SRS bands was used for morphology-, motility-, and texture-based features. The x-y centroid coordinates of each cell identified in TrackMate were used to connect these features back to the track ID.

### Longitudinal SRS microscopy reveals drug-induced alterations in live-cell proliferation dynamics

We applied our platform to investigate the temporal dynamics of A549 cells treated with paclitaxel (Pac), doxorubicin (Dox), lapatinib (Lap), osimertinib (Osi), or vehicle control (0.04% DMSO) over 24 hours (Figure 4A, Movies S2-6). Cells were treated at concentrations around their experimentally determined IC_50_ (Figure S3) for 24 hours. DMSO treated cells preserved a flattened, spindle-like morphology with stable membrane contours, indicative of unperturbed adhesion and cytoskeletal integrity. As shown in the zoom-in image Figure 4B, Pac-treated cells rapidly adopted a rounded, refractile appearance, consistent with microtubule stabilization and prolonged spindle-assembly checkpoint activation.^52^ Dox treatment induced pronounced membrane protrusions and cell shrinkage, hallmarks of DNA strand breaks and caspase mediated-apoptosis.^53^ Lap and Osi treatments produced partial cytoplasmic retraction and reduced spreading, reflecting blockade of EGFR/HER2 -driven mitogenic signaling pathways.^54, 55^ We also observed significant cell-cell adhesion with Osi treatment, creating the appearance of larger cells.

**Figure 4.**
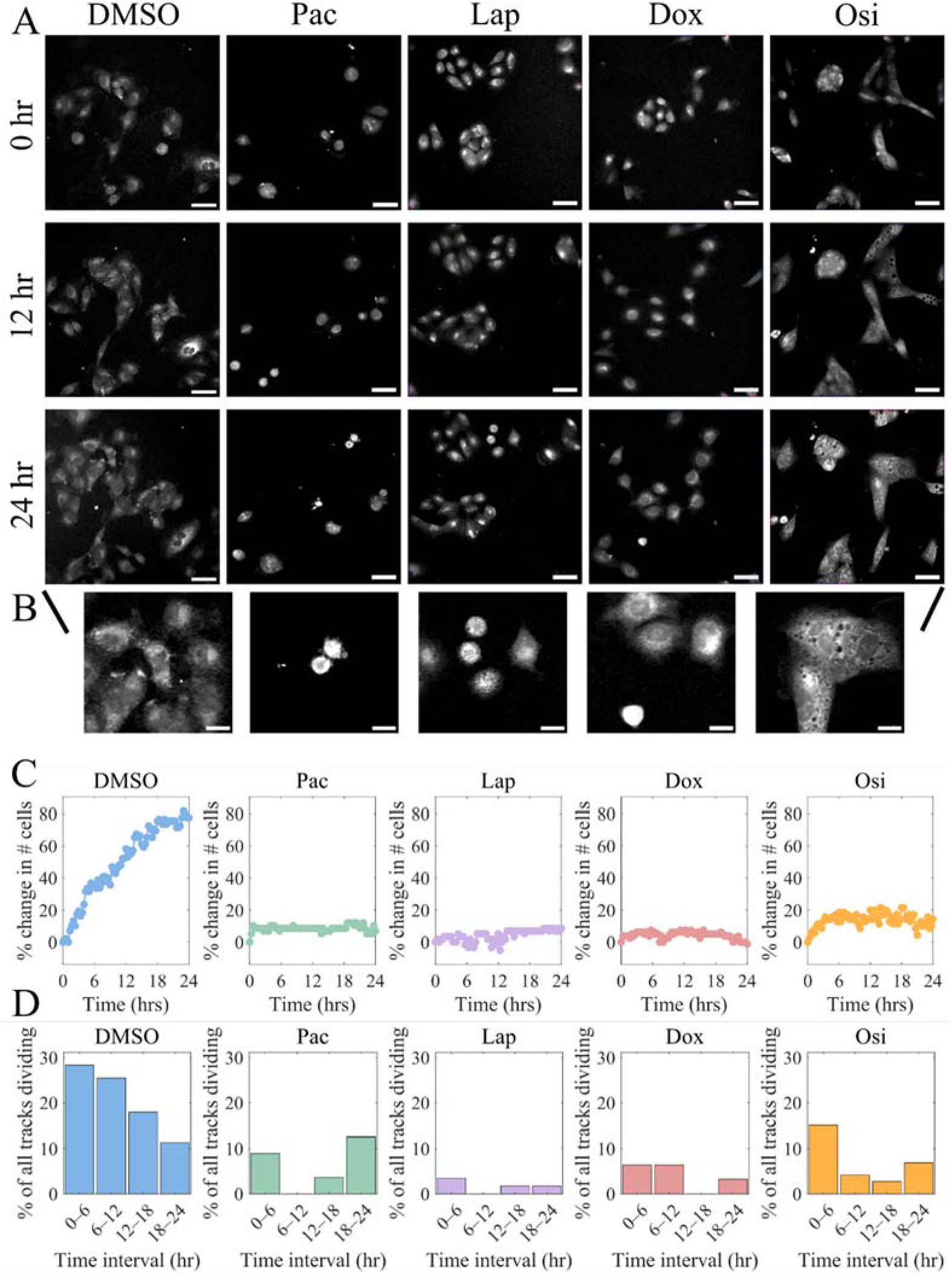
Dynamic cellular responses in drug-treated populations. A. Representative images of drug-treated cells over 24 hours of imaging. Each image is an intensity projection of lipid and protein SRS bands of one FOV of A549 cells treated with (left to right) 0.04% DMSO, 6 µM Osi, 10 µM Lap, 4 µM Dox, and 50 nM Pac. Scale bar: 50 µm. B. Zoomed in images at 24 hours. Scale bar: 20 µm. C. The number of tracked cells in each drug treatment condition over time. D. The percentage of mitotic events out of all tracked cells in each drug treatment condition represented in six-hour increments.

Quantitative analysis of cell proliferation revealed treatment-specific profiles over time (Figure 4C). Tracked DMSO cells showed ∼79 % net increase over 24 h. Comparatively, Pac and Dox limited net growth to <10 %, corroborating their potent antiproliferative and cytotoxic actions. Both Osi and Lap treatment also yielded pronounced cytostatic effect, likely due to the blockade of EGFR and downstream activities in EGFR wild-type A549 cells. ^54, 56^

Relatedly, analysis of mitotic entry timing (Figure 4D) revealed that in the DMSO-treated control population, the proportion of tracks undergoing mitosis peaked between 6–12 hr at approximately 35%, then continually declined stepwise at each time interval. This temporal decay in division frequency reflects intrinsic cell-cycle asynchrony. At the start of imaging, a subset of cells is ready to enter M phase and therefore drives the early mitotic wave. In contrast, their daughter cells and the remainder of the population must complete a full cell cycle before re-entering mitosis, yielding fewer divisions at later time points.^57^ Additionally, there can be confluency-dependent microenvironmental constraints, such as contact inhibition and localized growth factor depletion, that can delay subsequent mitotic entry. Compared to the control, all drug treated conditions show significantly less cell division. Intriguingly, both Pac and Osi exhibited a delayed rebound in divisions during the 18–24 hr window (Pac: ∼12%, Osi: ∼7 %). For Pac, this phenomenon aligns with mitotic slippage (Movie S7), wherein persistent spindle checkpoint arrest eventually yields cyclin B1 degradation and aberrant mitotic exit, often resulting in polyploid cells that may re-enter interphase.^52^ Osi’s late phase increase suggests partial inhibition of wild-type EGFR, resulting in an early suppression of mitotic entry comparable to Lap followed by a modest rebound to ∼7 % in the 18–24 h interval. This delayed recovery of division events likely reflects two factors: first, the reduced potency of Osi against wildtype EGFR necessitates higher drug exposure to maintain full cell cycle blockade; and second, a subset of cells may overcome sustained receptor inhibition and re-enter mitosis under prolonged exposure.^58–60^ When aggregated over 24 hr (Figure S4), average cumulative division rates were 76 % for DMSO, compared to 21% for Pac, 5% for Lap, 16 % for Dox, and 28 % for Osi. Collectively, these single-cell analyses demonstrate that drug action cannot be fully captured by endpoint assays alone and underscore the value of time-resolved imaging for uncovering how cells initially respond and subsequently adapt to pharmacological challenge.

### Spectral-morphological insight into drug-induced changes at the single-cell level

Next, we applied our platform to delineate the specific single-cell feature signatures that distinguish each drug-treated population. To do so, we utilized the extracted features from each single cell track and normalized the feature values against the average of the first three time points of each individual track. This creates a time-based feature change profile reflective of a drug’s mechanism of action. A few key features are represented in Figure 5 against the vehicle control (N=121).

**Figure 5.**
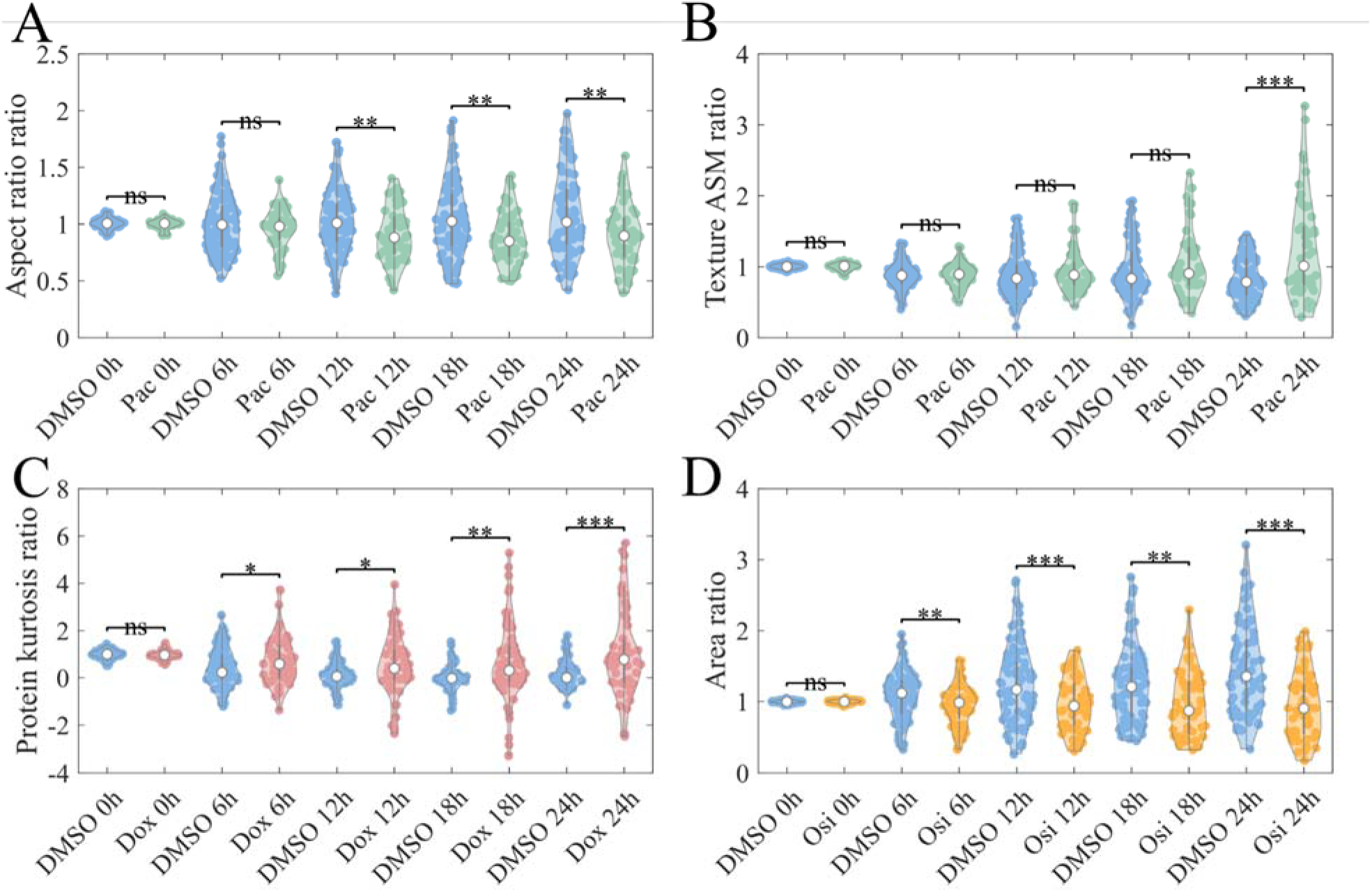
Selected features of phenotypic changes in drug-treated populations. Violin plots of single-cell feature ratios compared against DMSO-treated cells at six-hour time points. A. Aspect ratio changes in Pac-treated cell population. B. Texture angular second moment (ASM) ratio changes in Pac-treated cells. C. Protein kurtosis ratio changes in Dox-treated cells. D. Area ratio changes in Osi-treated cells.

For example, in Pac-treated cells (N=56) (Figure 5A), the aspect-ratio steadily decreases from DMSO control over time, indicating that cells are losing their elongated morphology and round as they are arrested at the spindle-assembly checkpoint.^52^ At the same time, it shows a large increasing spread in texture ASM ratio at 24 hr (Figure 5B), reflecting this mitotic arrest as a population of cells increase in texture uniformity, as illustrated in Figure 4B. With Dox treatment (N=94) (Figure 5C), protein kurtosis remains near baseline until ∼18 hr, then surges by 24 hr. Notably, there is a sharp spike in protein kurtosis around 1300 min, which appears to correlate with the formation of protein-rich apoptotic bodies before these dead cells are no longer tracked (Figure S5). Osi-treated A549 cells (N=73) showed a subtle continual decrease in cell area, suggesting cell aggregation.

As shown, our results highlight the ability of SRS microscopy to generate quantitative feature-ratio trajectories that serve as direct readouts of a compound’s mechanism of action with single-cell-precision. By capturing both the onset and temporal evolution of these phenotypic signatures, our approach can reveal not only the immediate efficacy of a drug but also heterogeneous and adaptive behaviors. Some drugs such as Lap (N=54) have less clear feature trends yet show cytostatic and motility effects only observed with longitudinal imaging. However, the averaging of features of all tracked cells in treatment conditions inherently buries information on single cell response. As all drug treatments were applied at approximately IC₅₀ doses, the resulting cell population is inherently heterogeneous: some cells continue to proliferate, while others undergo cell-cycle arrest or die. Single-cell tracking allows for differentiation between cell growth arrest and cell death, two parameters that are confounded in regular proliferation assays.^3^ SRS imaging has been shown to differentiate signatures of cell death pathways spectrally and morphologically,^61^ providing a label-free alternative to live/dead fluorescence assays. For example, living cells are characteristic by smooth, well spread contours, decently uniform texture, and high solidity (Figure S6, Movie S2, Movie S8). In contrast, a mitotic arrested cell (magenta, t=840 min) may show sharp features of elevated lipid and protein signal, highly regular boundaries and homogeneous texture, and reduced size. After death and fragmentation under Pac treatment (magenta, t=1320 min), there is a rise in heterogeneity and lipid signal, and a reduction in solidity. Overall, single cell features allow for detailed tracking of viability and processes of cell death. Due to the asynchronous nature of cell cycle and drug response, feature averaging introduces large variability that may obscure more subtle changes in morphological and chemical response of single cells.

### Tracing vacuole formation in drug-treated cell populations

Having observed single cell phenotypic trajectories, we then probed the occurrence of vacuolization made apparent in the Lap and Osi treatment conditions. We utilized our platform to observe this subcellular organelle remodelling, a hallmark of activated autophagic or endocytic processes.^62, 63^ By quantifying vacuole number, area fraction, and cumulative vacuole area in DMSO, Pac, Lap, Dox, and Osi treated A549 cells (Figure 6), we extend our timelapse SRS platform from whole cell feature profiling to organelle level phenotyping.

**Figure 6.**
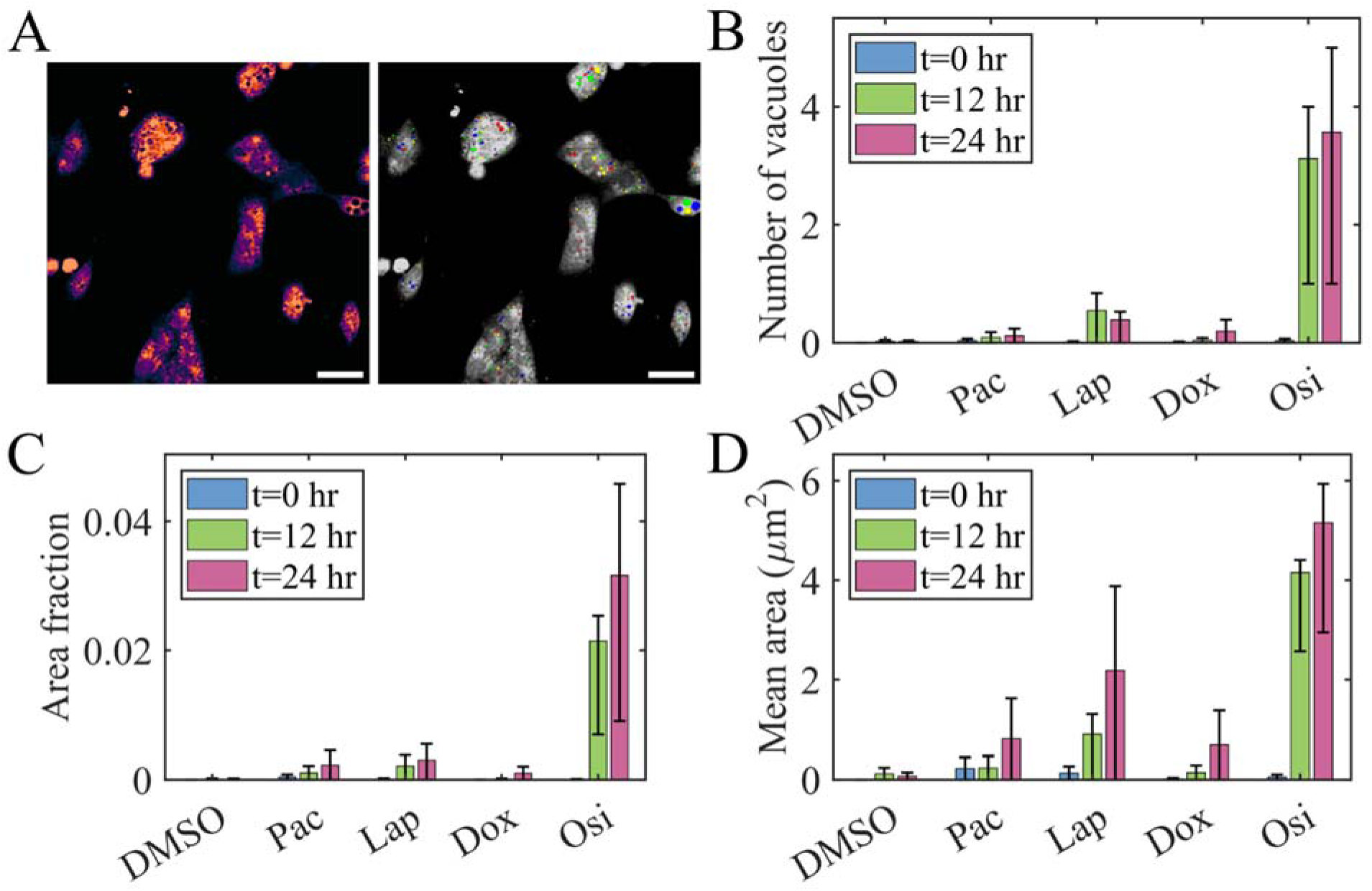
Vacuolization occurrence in drug treatment conditions. A. Vacuole segmentation (colored dots) from whole-cell image. B. Number of vacuoles (mean ± IQR) formed per cell in each drug-treated condition. C. Area fraction (mean ± IQR) of vacuoles per cell in each drug-treated condition. Mean vacuole area (mean ± IQR) per cell in each drug-treated condition.

In DMSO controls, vacuolization remains at basal levels (average <1 vesicle/cell; <0.01% area fraction, <1 µm^2^ mean vacuole area), reflecting housekeeping endolysosomal turnover. Pac and Dox produce only slight increases (average <1 vacuole/cell; <0.01% area fraction), likely indicating that neither prolonged mitotic arrest nor DNA damage at the tested doses engages autophagic vacuole biogenesis beyond baseline. In contrast, Lap elicits a moderate vacuolization response (average ∼0.38 vacuoles/cell by 24 hr; ∼0.03% area fraction; ∼2.1 µm^2^ vacuole area), consistent with receptor inhibition–induced autophagy, wherein EGFR/HER2 blockade activates AMPK and downregulates mTOR, promoting autophagosome formation.^64^ Strikingly, Osi triggers dramatic vacuole formation over time (average ∼3.5 vacuole/cell by 24 hr; ∼3% area fraction, ∼5.2 µm^2^ vacuole area), likely implying excessive autophagic flux under EGFR inhibition. These organelle level signatures illuminate how distinct drug classes diverge in their engagement of autophagy related and endocytic pathways.

### Single-cell migratory trajectories in drug-treated populations

Besides morphological features, we can leverage the dynamic tracking of single cells to extract cell motility features and evaluate single cell migratory behavior change in response to drug treatment. Figures 7A–E show single-cell trajectories under each treatment, with DMSO-treated cells exploring broadly (displacements >75 µm) while Pac, Lap, and Osi treatment markedly confine movement into tight space, reflecting suppressed exploratory motility. Dox-treated cells exhibit more extended early segments before retracting, hinting at an early mobilization followed by loss of persistence. In Figure 7F, the shape of the instantaneous speed curves makes these differences clearer: DMSO controls show a modest rise in speed through mid-course, followed by a decline, perhaps from crowding. Pac produces a moderate, relatively steady suppression of speed without dramatic early/late inflections. In contrast, the EGFR/HER2 inhibitors Lap and Osi suppress motility immediately and maintain a flattened, low-speed profile, consistent with sustained cytostatic damping of movement. Dox is distinct, as it shows an early higher speed that collapses later, likely suggesting an acute stress-related mobilization followed by pronounced loss of motility. The curve shapes reflect mechanistic differences in the drug types, showcasing how each uniquely reshapes cell dynamic behavior over time.

**Figure 7.**
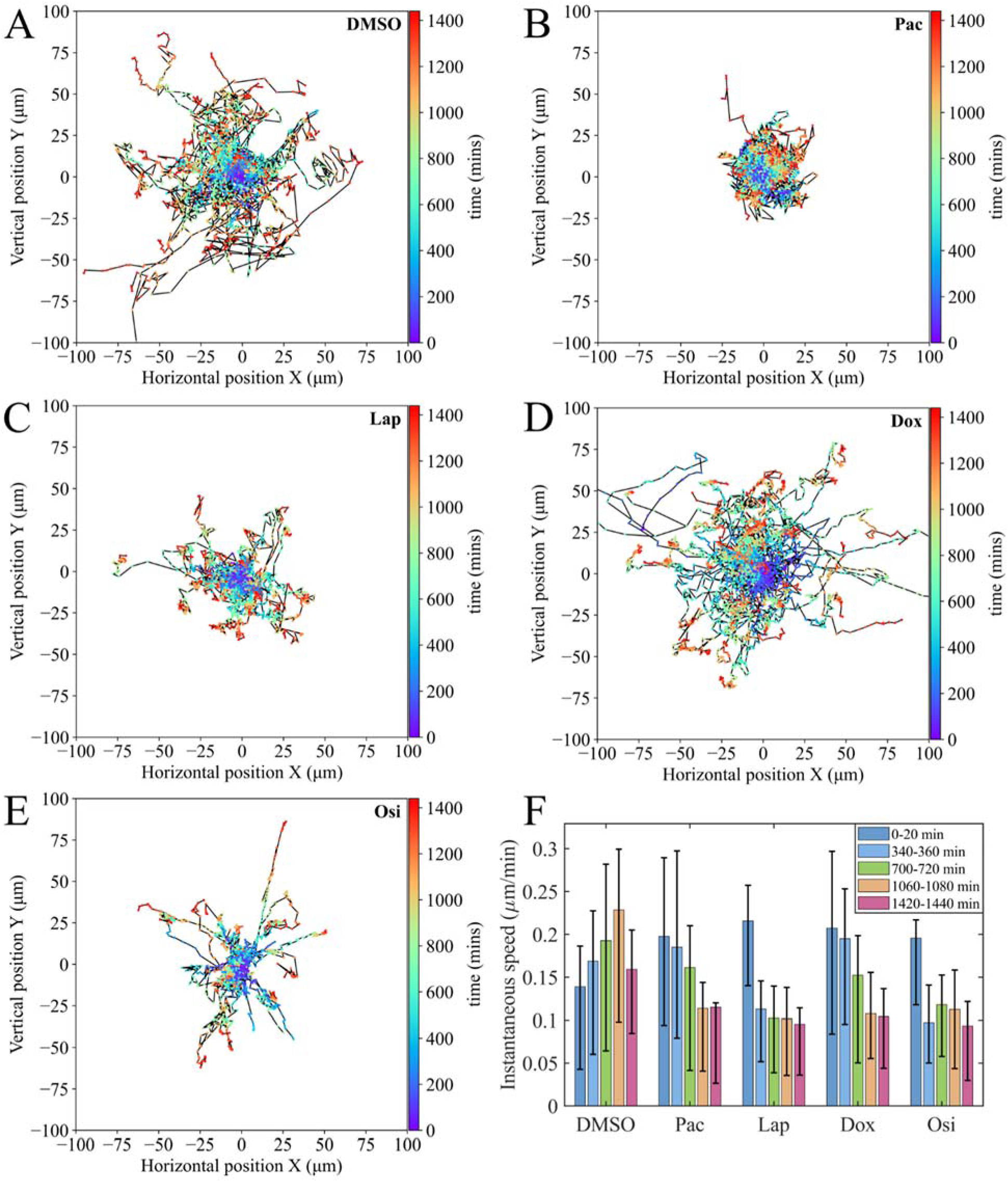
Drug-induced differences in cell motility. A-E. Single-cell trajectory plots (µm) where color bar indicates time and each line is one cell track. F. Instantaneous speed (µm/min) (mean ± interquartile range) calculated at different time intervals.

## Discussion

We have developed and validated a low cost, 3D-printed perfusion chamber tailored for label-free time-lapse SRS microscopy, enabling continuous drug delivery, environmental control, and high-resolution chemical imaging over 24 hr. By demonstrating equivalent fold change in cell counts under both static and perfused conditions, we confirm that our platform maintains physiological viability and minimizes photodamage during prolonged imaging. Leveraging this chamber, we implemented an automated image-processing pipeline comprising of single-cell segmentation and tracking across three SRS bands to extract multidimensional feature trajectories. Application to lung carcinoma cells treated with Pac, Dox, Lap, and Osi revealed mechanistically consistent dynamic phenotypes: microtubule destabilization and mitotic rounding under Pac; apoptotic blebbing with Dox; sustained cytostatic remodelling with Lap; and partial inhibition with late rebound and pronounced autolysosomal vacuolization under Osi. By extending our analysis from single cell morpho-chemical analysis to subcellular vacuole profiling and motility, we demonstrate the multidimensional resolving power of label-free SRS imaging. Feature-level signatures serve as quantitative metrics for drug mechanism, potentially allowing discrimination between cytotoxic, cytostatic, and adaptive stress responses at single-cell resolution.

Currently, our perfusion-imaging platform only maintains three independent flow channels, which constrains the number of conditions that can be tested simultaneously and raises the possibility of batch effects. The size of each well could be further reduced to increase the number of wells, at the expense of imaging time. Second, all experiments were conducted with a 20× objective to increase throughput and minimize photodamage, which limits spatial resolution and makes fully automated segmentation and tracking of densely packed cells challenging. In particular, the subtle membrane retractions and cell–cell adhesions seen in Osimertinib-treated cultures required extensive manual correction. Designing a version of our chamber compatible with a higher-NA objective would not only improve single-cell segmentation and tracking fidelity but also open the door to imaging smaller, organelle-level structures without sacrificing environmental control. Compared to the popular Cell Painting assay, our label-free SRS imaging still lacks the imaging throughput needed for large-scale drug screening. However, it has the distinct advantage of providing robust morphochemical information without relying on fluorescent staining, which not only obviates staining variability, but also broadens features beyond a few selected organelles.^65^

Looking forward, several extensions could dramatically expand the utility of perfusion-enabled SRS phenotyping. Advanced machine learning and deep learning algorithms could be used to extract single cell features that reliably predict cell fate and drug mechanism of action. Towards that end, computational synchronization of cell response could be a powerful approach to mitigate the intrinsic cell cycle and cell state variation and reveal divergent drug responses of subpopulation of cells. On the imaging side, by tuning the laser to fingerprint vibrations, we could directly measure intracellular drug uptake and distribution in real time, yielding pharmacodynamic maps at single-cell resolution. The platform is also well suited to more complex culture formats, co-cultures of tumor cells with stromal or immune cells, 3D spheroids or organoids, and even patient derived organoids, allowing us to probe how cell-cell and cell-matrix interactions modulate dynamic drug responses. Finally, scaling throughput to 24-well or even 96-well (or higher-density) formats would bring SRS-based screens into the realm of high-content drug discovery, marrying the chemical specificity of label-free imaging with the power of large-scale phenotypic screening.

## Author contributions

E.L.D.: Conceptualization; Methodology; Investigation; Formal analysis; Data curation; Writing – original draft; Writing – review & editing. K.C.: Formal analysis; Data curation; Software; Methodology. B.S.W.: Conceptualization; Methodology; Software. A.C.A.: Methodology. V.P.: Investigation. Y.W.: Methodology. D.F.: Conceptualization; Supervision; Writing – review & editing; Funding acquisition.

## Acknowledgements

This research was supported by the NIH R35 GM133435 to D.F.

