## supplemental figures and tables for "Label-Free Longitudinal Imaging of Single Cell Drug Response with a 3D-Printed Cell Culture Platform"

| Features Categories | Features | Input |
|---|---|---|
| Intensity statistics | 01. Total intensity<br>02. Mean intensity<br>03. Standard deviation<br>04. Skewness<br>05. Kurtosis<br>06. Minimum intensity<br>07. Max intensity<br>08. 10th Percentile<br>09. 25th Percentile<br>10. 50th Percentile<br>11. 75th Percentile<br>12. 90th Percentile | single channel image + cell mask |
| Morphology | 01. Area<br>02. Perimeter<br>03. Circularity<br>04. Roundness<br>05. Solidity<br>06. Extent<br>07. Bounding box area<br>08. Eccentricity<br>09. Major axis length<br>10. Minor axis length<br>11. Aspect ratio<br>12. Equivalent diameter<br>13. Orientation<br>14. Compactness<br>15. Maximum radius<br>16. Median radius<br>17. Mean radius<br>18. Feret diameter maximum<br>19. Feret diameter minimum<br>20. Zernike shape moments<br>21. Spatial moments<br>22. Central moments<br>23. Normalized moments<br>24. Hu moments<br>25. Inertia tensor elements and eigenvalues | Intensity-averaged image + cell mask |
| Texture | Haralick texture:<br>01. Angular Second Moment<br>02. Contrast<br>03. Correlation<br>04. Sum of Squares: Variance<br>05. Inverse Difference Momen<br>06. Sum Average<br>07. Sum Variance<br>08. Sum Entropy<br>09. Entropy<br>10. Difference Variance<br>11. Difference Entropy | Intensity-averaged image + cell mask |

| | | |
|---|---|---|
| | 12. Information Measure of Correlation 1<br>13. Information Measure of Correlation 2 | |
| Subcellular (vacuole statistics) | 01. Number of vacuole<br>02. Area Fraction<br>03. Sum of vacuole area<br>04. Min of vacuole area<br>05. Max of vacuole area<br>06. Mean of vacuole area<br>07. Standard deviation of vacuole area<br>08. Sum of vacuole perimeter<br>09. Min of vacuole perimeter<br>10. Max of vacuole perimeter<br>11. Mean of vacuole perimeter<br>12. Standard deviation of vacuole perimeter | cell mask + vacuole mask |

**Supplemental Table 1.** List of features extracted from SRS cell images.

| Item | Vendor | Part Number / Model | Quantity per device | Unit Price |
|---|---|---|---|---|
| **ITO glass slide** | Delta Technologies | CG-81IN-0115 (25×75×1.1 mm), 30–60 Ω | 1 | $3.09 each |
| **FEP-lined Versilon PVC tubing** | McMaster-Carr | Ultra-Chemical-Resistant Clear Versilon PVC Tubing 1/16″ ID × 1/8″ OD 50 ft | 1 | ≈ $2.86 per ft / $143 per 50 ft |
| **Stainless steel tubing** | McMaster-Carr | Miniature stainless steel tubing with 90° bend / 5560K81 | 8 | $5.61 each / $44.88 |
| **NTC thermistor** | Newark Electronics | NTCLE100E3103JB0 / 10 kΩ | 1 | $0.33 |
| **PID temperature controller** | Omega Engineering (from Newark Electronics) | CN38S-TC-R1-R2-12V | 1 | $277.86 |
| **Peristaltic pump** | Longer Precision Pump Co. | L100-1S-2 (8 channel) | 1 | From $916.00 |
| **PDMS kit** | Dow Chemical Company (from Electron Microscopy Sciences) | SYLGARD 184 Silicone Elastomer kit, 1.1 kg | 1 | $180 per kit |
| **PLA filament** | Bambu Lab | PLA Basic; 1 kg spool | 11.62 g | $0.27 / $22.99 for 1 kg |

**Supplemental Table 2.** Cost of supplies for 3D-printed perfusion chamber.

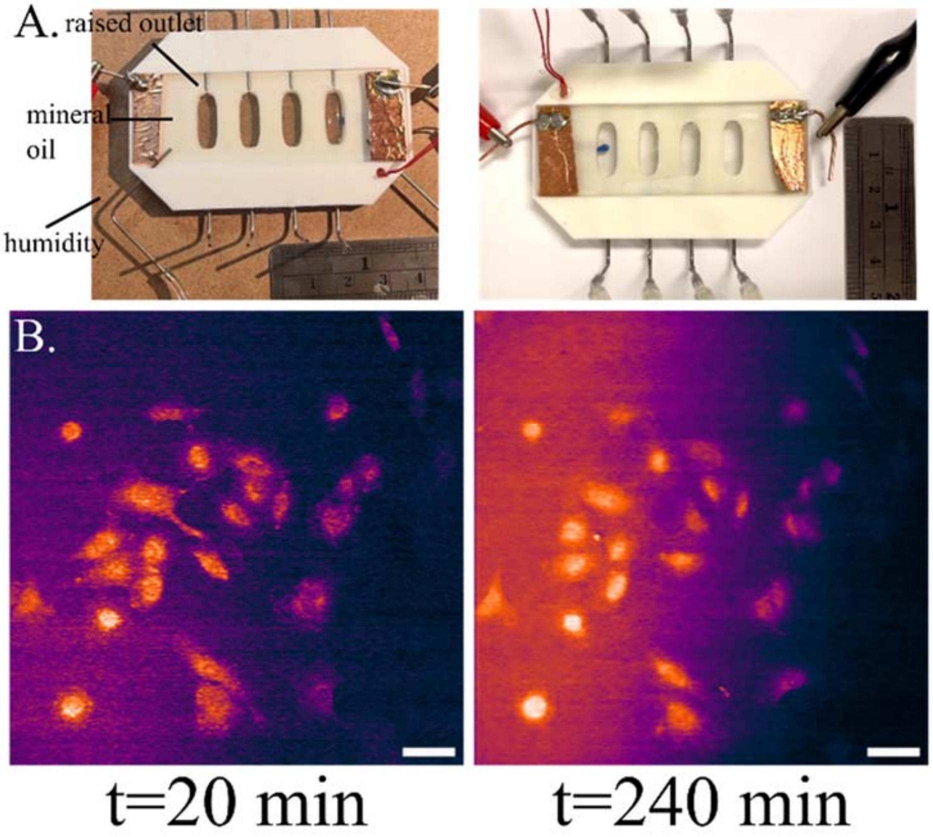


**Supplemental Figure 1.** A. Comparison of open dish (left) versus closed dish design (right) with indicated differences. B. Longitudinal SRS imaging (average water and protein band) using an open 3D-printed perfusion chamber design with mineral oil. Scale bar: 20 µm.

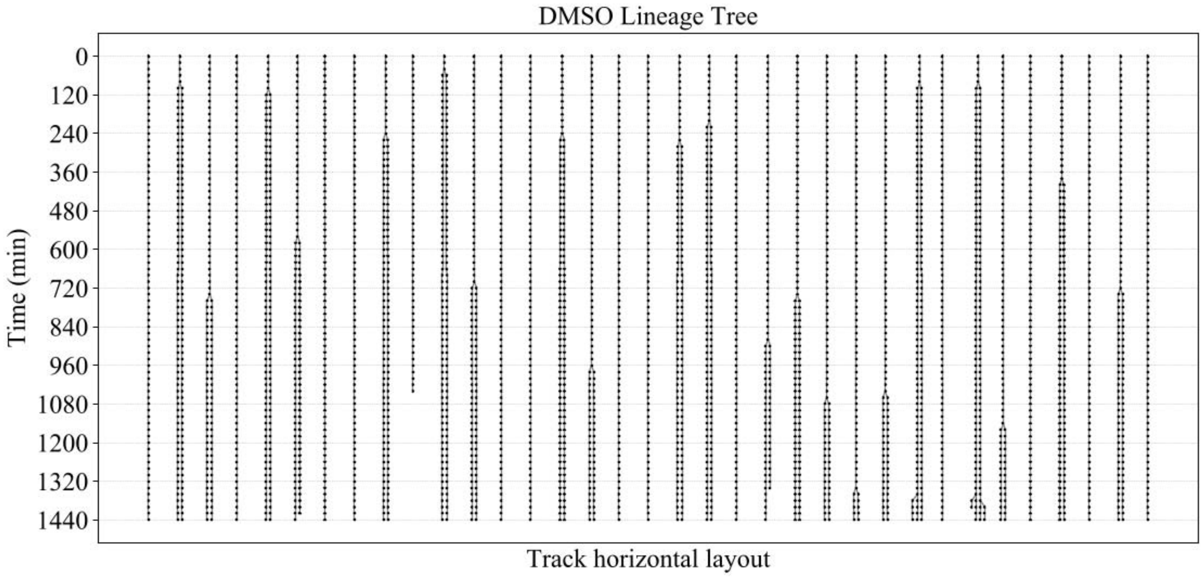


**Supplemental Figure 2.** Lineage-tree reconstruction from single-cell tracking.

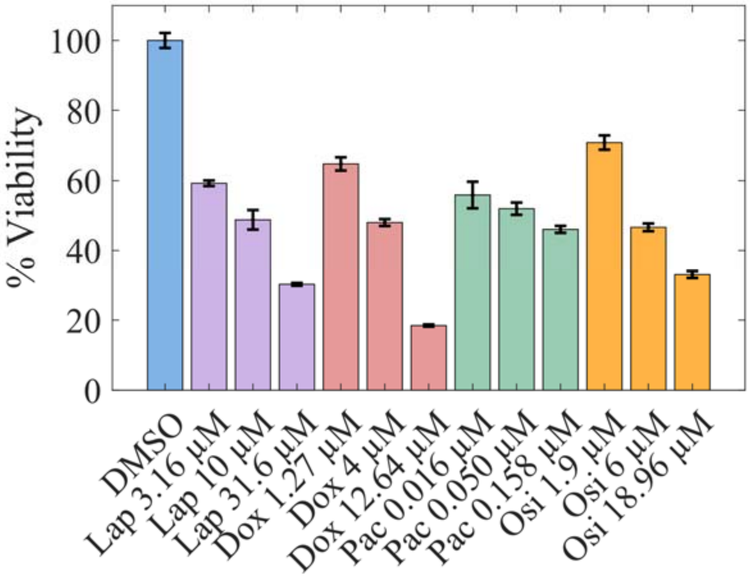


**Supplemental Figure 3.** MTS viability assay on drug-treated A549 cells. A549 cells were treated for 24 hours at ½-log scale concentrations. N=4 for each condition.

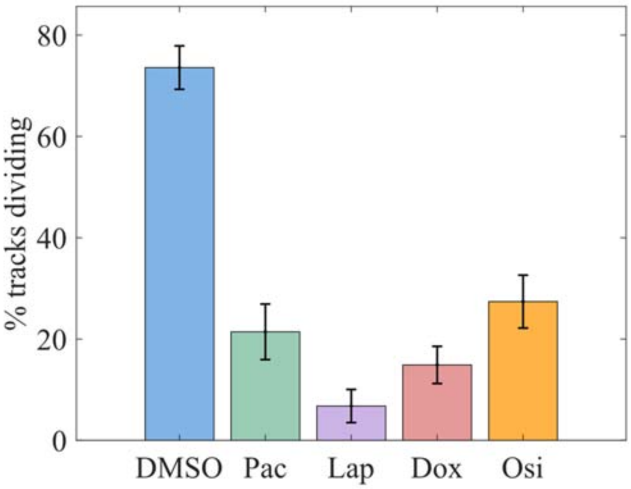


**Supplemental Figure 4.** Percentage of all cell tracks that undergo a mitotic event.

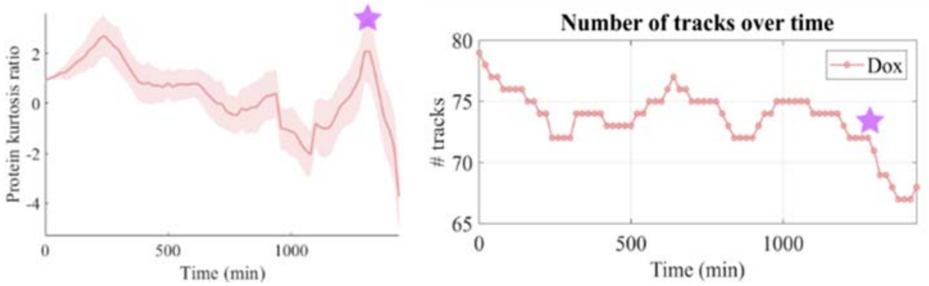


**Supplemental Figure 5.** Number of cell tracks under Dox treatment. Starred: moment of large decrease in cell count correlative with an increase in protein kurtosis.

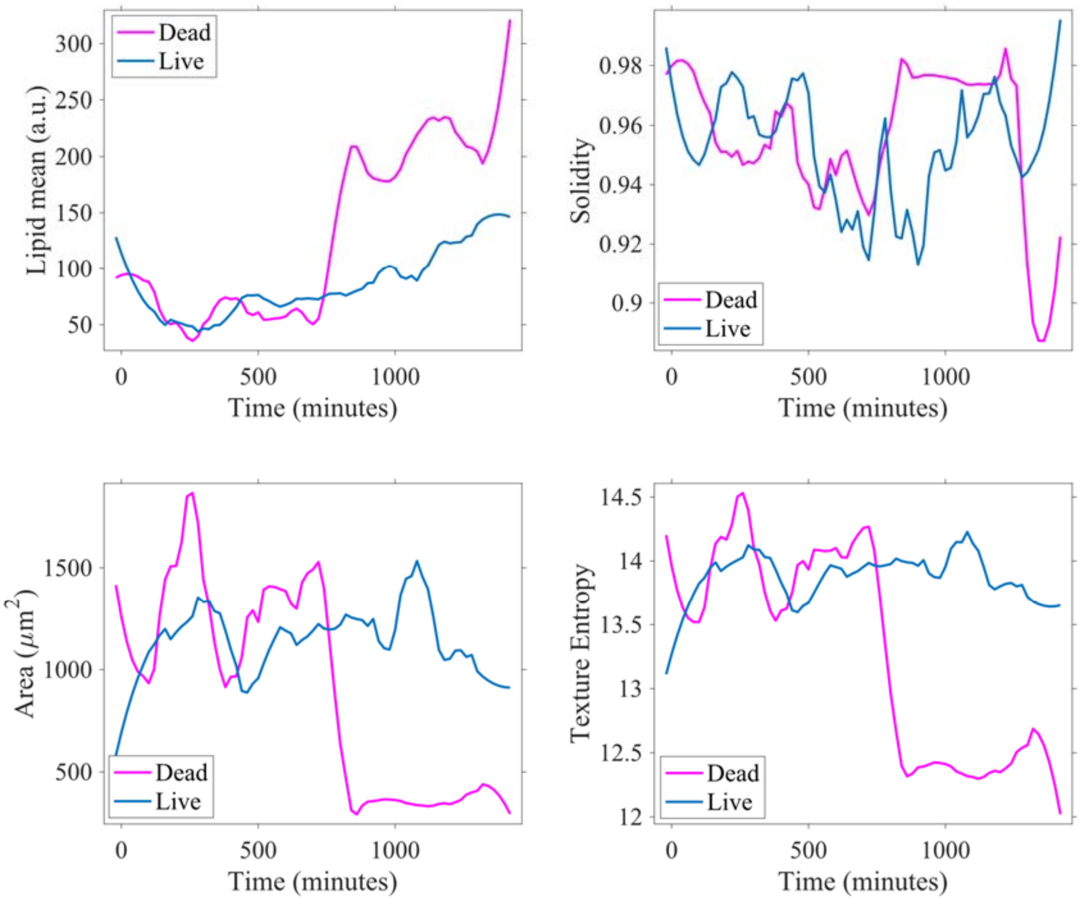


**Supplemental Figure 6.** Temporal dynamics of features of a living (blue) and dying (magenta) cell in Pac-treated cell population.
